# Epigenetic editing at individual age-associated CpGs affects the genome-wide epigenetic aging landscape

**DOI:** 10.1101/2024.06.04.597161

**Authors:** Sven Liesenfelder, Mohamed H. Elsafi Mabrouk, Jessica Iliescu, Monica Varona Baranda, Athanasia Mizi, Martina Wessiepe, Argyris Papantonis, Wolfgang Wagner

## Abstract

Aging is reflected by genome-wide DNA methylation changes, but it is largely unclear how these epigenetic modifications are regulated. In this study, we explored the possibility to interfere with epigenetic clocks by epigenetic editing at individual CpG sites. CRISPR-guided approaches (dCas9-DNMT3A and CRISPRoff) facilitated targeted methylation at an age-associated genomic region in *PDE4C* that remained stable for more than three months. Furthermore, epigenetic editing evoked many genome-wide off-target effects, which were highly reproducible and enriched at other age-associated CpGs – thus, they are not random off-target effects, but seem to resemble coregulated epigenetic bystander modifications. 4C chromatin conformation analysis at age-associated sites revealed increased interactions with bystander modifications and other age-associated CpG sites. Subsequently, we multiplexed epigenetic modifications in HEK293T and primary T cells at five genomic regions that become either hypermethylated or hypomethylated upon aging. While epigenetic editing at age-hypomethylated CpGs appeared less stable, it also resulted in a clear enrichment of bystander modifications at other age-associated CpGs. Conversely, epigenetic clocks tend to be accelerated up to ten years after targeted DNA methylation, particularly at hypermethylated CpGs. These results demonstrate that targeted epigenome editing can modulate the epigenetic aging network in its entirety and thereby interfere with epigenetic clocks.

## Introduction

The epigenetic landscape changes continuously during human aging. This is particularly reflected by increased or decreased DNA methylation (DNAm) at specific CpG dinucleotides in the genome ^1^. Due to the steady and reproducible nature of these changes epigenetic clocks are widely used as a biomarker for aging ^2,3^. Accelerated epigenetic age is associated with higher all-cause mortality, indicating that these epigenetic signatures rather reflect biological than chronological age ^4-7^. While age-associated DNAm changes are well characterized as a predictor, it is largely unclear how they are coherently modified across the genome and if they directly affect the aging process ^3,8^. A better understanding of the underlying molecular mechanisms might ultimately allow control of age-associated epigenetic changes and thereby eventually even of the aging process.

So far, the methods to reset the epigenetic clocks are limited. It is possible to rejuvenate cells by reprogramming them into induced pluripotent stem cells, but this involves a complete resetting of epigenetic and functional characteristics of the cells ^9,10^. It is yet unclear if partial reprogramming can stably interfere with epigenetic clocks, while maintaining cellular identity ^11^. Demethylating agents or knockout of DNA methyltransferases (DNMTs) may also interfere with epigenetic clocks, but they do not control the complex and site-specific gains and losses of DNAm during aging. More recently, epigenetic editing approaches have been developed that facilitate site-specific modification of the DNAm pattern. For instance, the CRISPR toolbox uses single guide RNAs to direct a fusion protein of the nuclease deficient Cas9 with DNMT3A/3L (dCas9-DNMT3A) to specific sites in the genome ^12^. While DNMT3A-mediated epigenome editing is rather transient at most regions and usually lost within a few days ^13^, more complex fusion proteins have been developed for more sustained manipulations. For example, the “CRISPRoff” includes a transcriptionally repressive Krüppel-associated box (KRAB) domain in addition to DNMT3A/3L. KRAB recruits TRIM28/HP1α and thus increases the stability of DNA-hypermethylation to provide a suitable tool for gene silencing ^14-16^. However, it has been shown that dCas9-methyltransferases can have a surprisingly ubiquitous nuclear activity with many potential off-target effects ^17^.

So far, the epigenetic editing has not been used systematically to interfere with epigenetic clocks - and this approach may seem counterintuitive, given that aging is reflected by genome-wide changes in the DNAm pattern, rather than at individual CpG sites. On the other hand, there may be crosstalk of epigenetic modifications within a network that are yet to be elucidated. For example, we have recently observed in acute myeloid leukemia (AML) that patient-specific aberrant DNAm patterns were always symmetric on both alleles, which may be governed by an inter-allelic epigenetic crosstalk ^18^. Furthermore, age-associated DNAm was shown to evolve coherently resulting in heterogeneity of epigenetically younger to older cells in single cell analysis ^19,20^. Some methylation changes seem to originate from CpG clusters are coherently modified with age ^21^. To further investigate the potential co-regulation of age-associated DNAm in a network-like manner, we conducted epigenetic editing at epigenetic clock sites. Our findings provide evidence that local changes in the DNAm can result in a genome-wide response of aging signatures, thereby interfering with epigenetic clocks.

## Results

### CRISPR-guided epigenetic editing is site specific and stable

Initially, we aimed to modify DNAm in a genomic region within phosphodiesterase 4C (*PDE4C)*, which was identified as one of the first sites to gain age-associated DNAm across different tissues ^1^ and has a very high correlation with chronological age in blood ^10,22^. We used two alternative constructs to modulate this region in HEK293T cells: dCAS9-DNMT3A ^12^ and CRISPRoff ^14^ (both n = 3; Figure 1a). Pyrosequencing validated a highly significant increase in DNAm across all seven CpGs in this amplicon (P < 0.001; n = 3; Figure S1a). Furthermore, Illumina BeadChip (EPIC) analysis 14 days after transfection showed the highest gain of DNAm at the target site in *PDE4C* (Figure 1b). Both analysis methods indicated that up to 40% of the DNA strands in *PDE4C* have gained DNAm.

**Figure 1:**
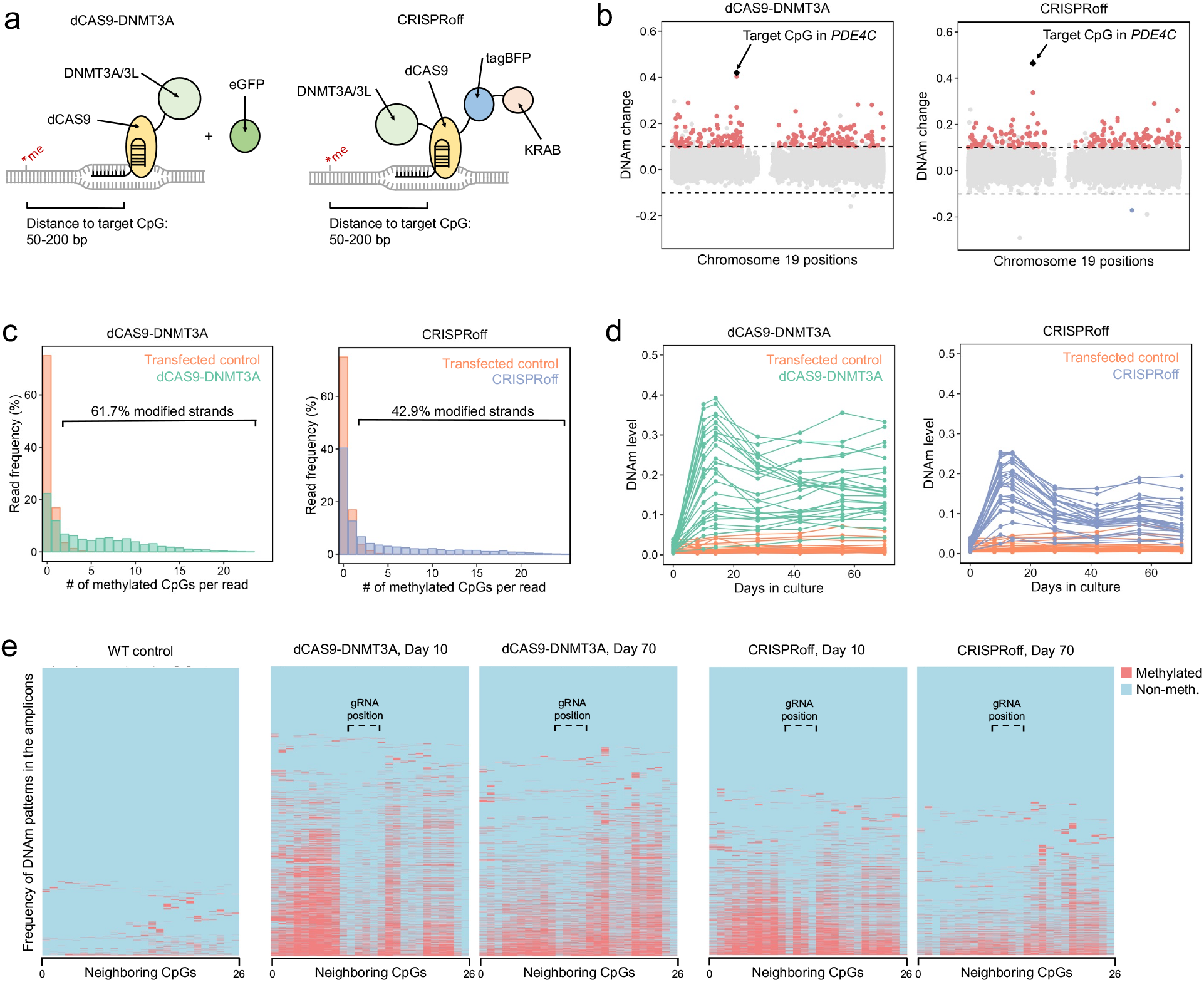
Epigenetic editing is stable but not coherent within the target region. **a)** Schematic presentation of two CRISPR-guided epigenetic editors used in this study. Deficient CAS9 protein is linked to DNMT3A/3L at the c-terminus (dCAS9-DNMT3A/3L, co-expressed with eGFP for selection) ^12^, and CRISPRoff comprising DNMT3A/3L at the n-terminus, and tagBFP and a KRAB domain at the c-terminus ^14^. Single guide RNAs were designed 50 to 200 base pairs distant of the target CpG. **b)** Manhattan plot illustrating the DNAm changes upon targeting the age-associated hypermethylated region in *PDE4C* (EPIC Illumina BeadChip data of chromosome 19; mean of three replica). Significant DNAm changes (delta mean DNAm > 0.1 and an adj. p-value < 0.05) are highlighted in red. The target CpG in *PDE4C* is highlighted in black. **c)** Bisulfite amplicon sequencing of 26 neighboring CpGs at the target region of *PDE4C*. The bar plot depicts the frequency of methylated CpGs on individual reads, indicating that even on modified DNA strands not all neighboring CpGs become coherently methylated. 61.7% and 42.9% of reads have higher methylation levels then observed in the wildtype, for dCAS9-DNMT3A and CRISPRoff, respectively. **d)**Time-course experiment of DNAm at *PDE4C* measured by bisulfite barcoded amplicon sequencing. The lines resemble the 26 different CpGs in the amplicons. **e)**Frequency of DNAm patterns in bisulfite barcoded amplicon sequencing data of *PDE4C*. Reads are clustered by their DNAm pattern. The binding region of one gRNAs is indicated. The second gRNA binds two basepairs next to the CpG #26 and might explain the low methylation gain at #25 and #26.

To get better insight into the DNAm changes at this region, we performed bisulfite amplicon sequencing that covered 26 neighboring CpGs. We anticipated that the fraction of modified DNA strands might be even higher than reflected by the mean DNAm at individual CpGs. To this end, we analyzed the frequency of methylated CpGs in individual reads: we hardly observed two or more methylated CpGs in the controls, whereas 61.7% and 42.9% of reads showed higher methylation levels in dCAS9-DNMT3A and CRISPRoff modified cells, respectively (Figure 1c). This demonstrates that due to heterogeneity of DNAm at neighboring CpGs epigenetic editing was even more efficient than indicated by crude DNAm levels at individual sites. Furthermore, we could demonstrate hypermethylation was stable over at least 100 days for both dCAS-DNMT3A and CRISPRoff – even though the cells were highly proliferative and lost the transiently transfected plasmids within few days (Figure 1d). This was also confirmed by pyrosequencing (Figure S1b).

Initially we expected that when a CRISPR-DNMT3A construct is targeted to a specific genomic region, all neighboring CpGs should be coherently modified. However, as mentioned above, the bisulfite barcoded amplicon sequencing data demonstrated that modifications at the 26 neighboring CpGs in the amplicon were overall not coherently modified (Figure S1c). Particularly with the dCAS9-DNMT3A construct, we observed less gain of DNAm at guide RNA binding position (Figure 1e). Notably, during the time course, this lowly methylated region assimilated towards the higher DNAm at neighboring CpGs with time (Figure S1d). This indicates local DNAm levels within the target region become more homogeneous over several months, albeit the transient CRISPR-guided DNA methyltransferases are no longer present.

### Epigenetic editing evokes highly reproducible genome-wide bystander modifications

While the highest gains of DNAm were observed at the target site, there were also many off-target effects. When we analyzed EPIC BeadChip data of the three corresponding replica, we observed significant increases of DNAm (abs. diff. beta-value > 0.1 and p-value < 0.05) at 4864 sites and 3326 sites for dCAS9-DNMT3A and CRISPRoff, respectively (Figure 2a,b; Figure S2a,b). Furthermore, there was even a significant decrease in DNAm at either 7 or 14 CpGs upon epigenome editing with these methods, which is in line with previous observations ^23^ and may not be explained by the activity of the constructs. Direct comparison of DNAm changes with dCAS9-DNMT3A and CRISPRoff revealed a correlation of R^2^ = 0.53 for all CpGs, indicating that the gains of methylation are not just random footprints of the constructs, but rather consistent epigenetic bystander modifications (Figure 2c). Notably, even the hypomethylation was rather consistent with both editors.

**Figure 2:**
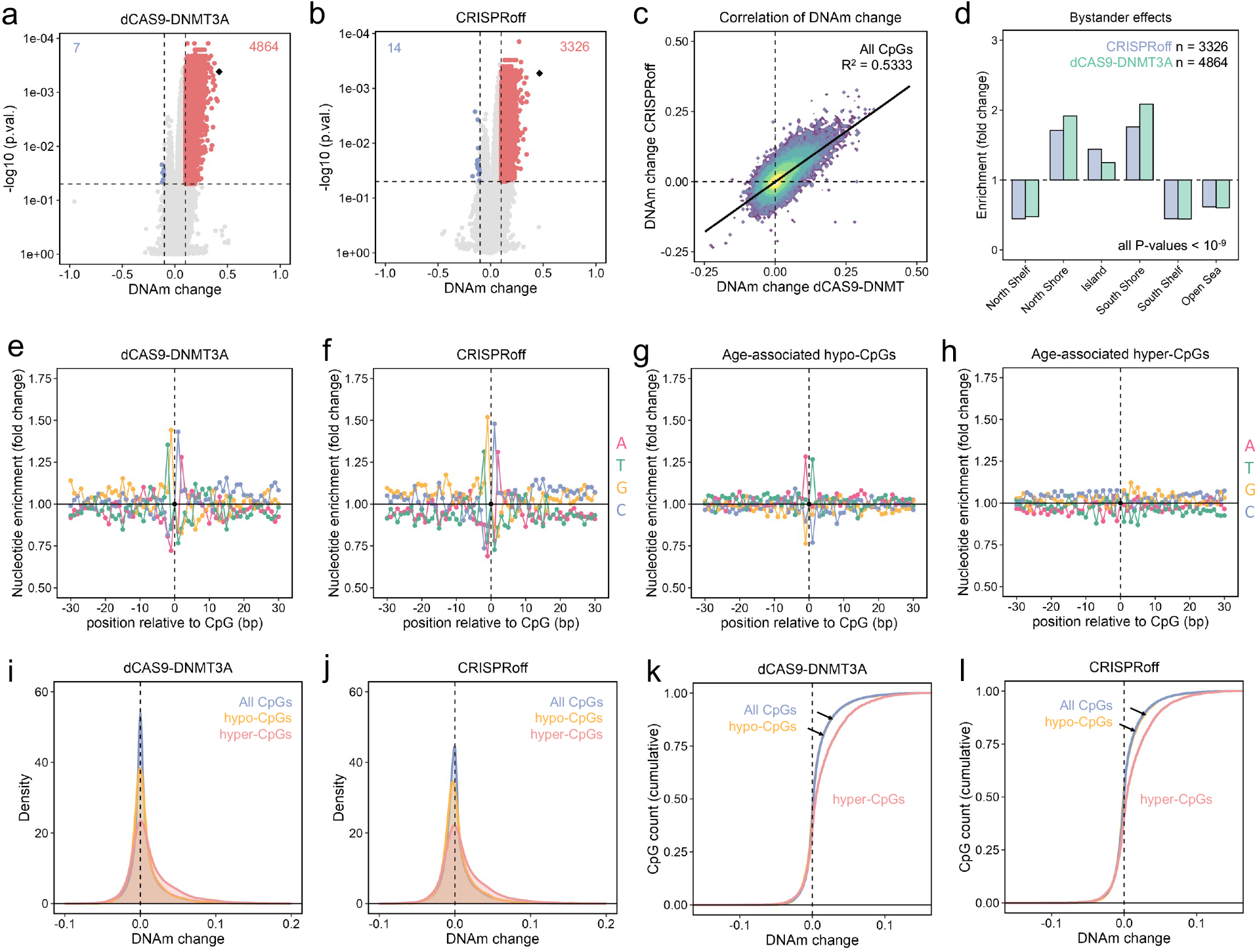
Genome-wide bystander effects of epigenome editing. **a**,**b)** Significant DNAm changes upon epigenetic editing at *PDE4C* with either dCas9-DNMT3A (a), or CRISPRoff (b). Vulcano plot depict significantly hypermethated (red) and hypomethylated (blue) CpGs and their numbers are indicated (n = 3). **c)**Scatterplot comparing DNAm changes in the dCAS9-DNMT3A and CRISPRoff experiments. Pearson correlation (R^2^ = 0.53) indicates that there is a high reproducibility of the bystander modifications, even with the different epigenetic modification approaches. **d)**Enrichment of the bystander effects in relation to CpG islands (CGI), and at shore and shelf regions surrounding CGIs. Enrichment was calculated in relation to all CpGs on the array and was highly significant for all categories (P < 10^-9^, Chi^2^ test). **e**,**f)** Relative frequency of nucleotides next to CpGs with epigenetic bystander modificaitons normalized to the entirety of CpGs on the BeadChip. Guanin and cytosine were overrepresented at the -1 and +1 flank positions, whereas thymine and adenine were enriched at the -2 and +2 positions. **g**,**h)** Relative frequency of nucleotides next to CpGs that become either hypomethylated with age (g; 4389 CpGs), or hypermethylated with age (h; 5328 CpGs) in a large-scale epigenome-wide association study ^25^. Adenine and thymidine were overrepresented in the -1 and +1 flank positions of hypomethylated CpGs. **i**,**j)** Distribution of epigenetic bystander modifications in the dCAS9-DNMT3A experiments (i) and the CRISPRoff experiments (j) was analyzed for all CpGs on the array, for the 4389 CpGs with age-associated hypomethylation, and 5328 CpGs with age-associated hypermethylation ^25^. Age-associatiate hypermethylation was enriched in CpGs that gain DNAm upon epigenetic editing. **k**,**l)** Cumulative distribution of the density functions in (i,j) to better visualize that bystander effects upon targeting *PDE4C* are enriched at other age-hypermethylated CpGs.

Subsequently, we tested if the bystander-effects were enriched at genomic regions that were predicted to have potential CRISPR-mediated off-target effects. Based on sequence homologies to our guide RNAs 123 regions were predicted to be potential off-target sites, but we did not observe any methylation changes here (Figure S2c,d). Instead, hypermethylated bystanders were ubiquitously enriched in islands and shores, but decreased in the shelf and open sea (P < 10^-9^; Figure 2d). Furthermore, the bystander effects were rather observed in upstream promotor regions (Figure S2e). We further analyzed the frequency of nucleotides at neighboring positions of bystander effects. Indeed, bystanders of epigenome editing featured distinct nucleotides around the differentially methylated CpGs: Guanin and cytosine were present at the -1 and +1 position about 41-52% more often than for random CpGs. Besides, thymine and adenine were enriched at the -2 and +2 position, respectively (Figure 2e,f). This nucleotide pattern around bystander CpGs has moderate association with the flank sequence preferences of the enzyme DNMT3A ^24^.

### Bystander modifications are enriched at other age-associated CpGs

Next, we sought to explore if the epigenetic bystander modifications upon targeting *PDE4C* are potentially related to other epigenetic clock sites. To this end, we utilized results of a large-scale epigenome wide association study of 18,413 individuals that identified the top 10,000 CpGs linearly associated with age ^25^. Of these, 5328 hypermethylated and 4389 hypomethylated CpGs passed our filter criteria for CpGs to be analyzed. We did not observe a clear motive for age-associated hypermethylation, whereas the hypomethylated CpGs showed a complementary enrichment of adenine and thymine at the -1 and +1 position, which was in line with previous reports ^26^ (Figure 2g,h).

Subsequently, we tested if the bystander DNAm changes upon modification of *PDE4C* were enriched in age-associated hyper- or hypomethylated CpGs. Strikingly, the 5328 CpGs with age-associated hypermethylation showed a significant methylation gain compared to all other or random CpGs – and this was observed for modification with either dCAS9-DNMT3A or CRISPRoff (P < 2.2e-16, Figure 2i-l). This indicates that the epigenetic bystander modifications of *PDE4C* are associated with other age-associated DNAm changes.

### Chromatin interactions contribute to bystander effects

To better understand why the bystander modifications occur in this highly reproducible manner, we investigated transposase-accessible chromatin with high-throughput sequencing (ATAC-seq) data of HEK293T cells ^27^. In fact, the epigenetic bystander modifications upon targeting *PDE4C* with either dCAS9-DNMT3A or CRISPRoff were clearly enriched at open chromatin (Figure 3a, P < 10^-15^). In contrast, CpGs with age-associated hypomethylation were rather associated with closed chromatin (Figure 3b, P < 10^-15^). Conversely, age-associated hyper-CpGs tended to have higher chromatin accessibility (P < 10^-15^). Thus, at least part of the bystander modifications might be attributed to chromatin accessibility. However, performing logistic regression analysis on methylation changes and chromatin acessibilty this association was less evident, indicating that chromatin state alone does not sufficiently explain the bystander effects with epigenome editing (Figure S3a, R^2^ = 0.0689 for hyper- and R^2^ = 0.0572 for hypo-CpGs).

**Figure 3:**
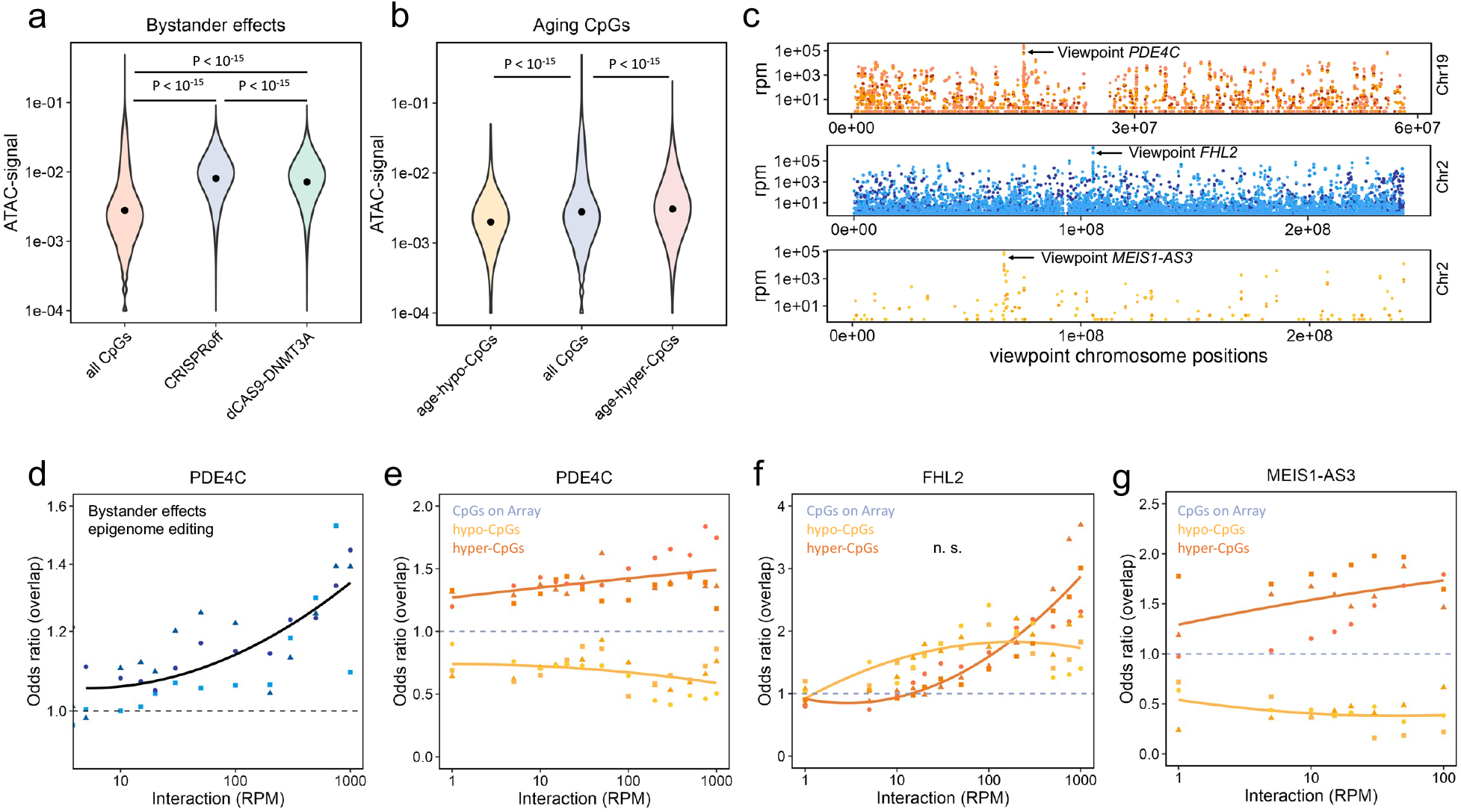
Chromatin conformation contributes to epigenetic bystander modifications. **a)**ATAC-seq data of HEK293T cells ^27^ was used to estimate chromatin accessibility at epigenetic bystander modifications upon modification of *PDE4C* with either dCas9-DNMT3A or CRISPRoff. The higher ATAC-seq signals indicate that bystander modifications are enriched at open chromatin regions. **b)**In analogy, we tested if CpGs with age-associated DNAm changes are also associated to chromatin accessibility in the ATAC-seq data HEK293T cells. The 4389 age-hypomethylated CpGs revealed overall lower ATAC signal, whereas the 5328 age-hypermethylated CpGs were enriched at open chromatin. **c)**Intrinsic 4C-sequencing in HEK293T cells for three age associated sites in *PDE4C, FHL2*, and *MEIS1-AS3* (triplicates per viewpoint). The Manhattan plots depict the number of reads (rpm = reads-per-million) at the cis-interacting regions on the corresponding chromosomes. The highest signal was observed around the 4C viewponts. **d)**Bystander effects become enriched with increasing coverage in 4C-sequencing. The association is best described by an exponential model (n = 3, different symbols correspond to replica; R^2^: 0.58, p-value < 10^-15^). **e-g)** Overlap of 4389 age-hypo and 5328 age-hypermethylated CpGs at interacting genomic regions in the 4C-sequencing data for *PDE4C* (e), *FHL2* (f), and *MEIS1-AS3* (g).

Next, we investigated if the coherent modification of bystander DNAm changes might also be attributed to the higher order of chromatin. Therefore, we conducted intrinsic 4C-sequencing to identify chromatin that interacted with the target site in *PDE4C*. For comparison, we considered an age-associated hypermethylated region in Four and a half LIM domains protein 2 (*FHL2*) and a hypomethylated region in MEIS1 antisense RNA 3 (*MEIS1-AS3;* all n = 3, Figure S3b,c). As expected, signals with the highest reads-per-million (rpm) counts were localized close to the viewpoint (Figure 3c). Excluding the *PDE4C* target region from analyses, we observed that bystander modifications (abs. diff. beta-value > 0.1 and p-value < 0.05) were increasing with higher interaction to *PDE4C* (Figure 3d). This association was best described by exponential modelling for the highly interacting CpGs (residual SE = 0.076, R^2^ = 0.53, P < 10^-15^). Therefore, at least some of the bystander modifications could be attributed to interacting chromatin.

To explore whether age-associated genomic regions might be generally enriched in interacting chromatin, we tested the enrichment of the 5328 hyper- and 4389 hypomethylated CpGs at the interaction of the three viewpoints. In fact, particularly the CpGs that become hypermethylated with aging were enriched in the interactome of all three age-associated CpGs (Figure 3e-g). Considering that *PDE4C* and *FHL2* revealed larger interactomes as compared to *MEIS1-AS3* (Figure S3b,c), it appears that particular CpGs with age-associated hypermethylation seem to have chromatin interaction with each other. Overall, the spatial interaction of epigenetic bystander modifications and of age-associated CpGs further pointed towards an interacting epigenetic network.

### Multiplexed epigenetic editing at age-associated CpGs

We reasoned that targeting multiple age-associated CpGs might increase interferences with a possible epigenetic aging network. Therefore, we multiplexed targeting of five age-associated regions: *PDE4C, FLH2, ELOVL2, KLF14*, and *TEAD1* ^22,28^. These experiments were again performed in HEK293T cells with a scramble guide RNA for control (n = 3; Figure 4a). Seven days after transfection, EPIC Illumina BeadChip analysis demonstrated clear gains of DNAm particularly at regions in *PDE4C, FHL2*, and *KLF14* (Figure 4b). These changes were not restricted to the target CpG, but enriched in a 1.5 kb area (Figure 4c). The gain of DNAm was further validated by pyrosequencing and remained stable at the age-associated sites for at least 17 days (Figure S4) – however the site-specific gain of DNAm was less pronounced and more variable than observed with the target site *PDE4C* alone. This might also be a reason why we did not observe significant bystander modifications at other genomic regions and hence we did not observe a clear enrichment at age-associated regions (Figure 4d,e).

**Figure 4:**
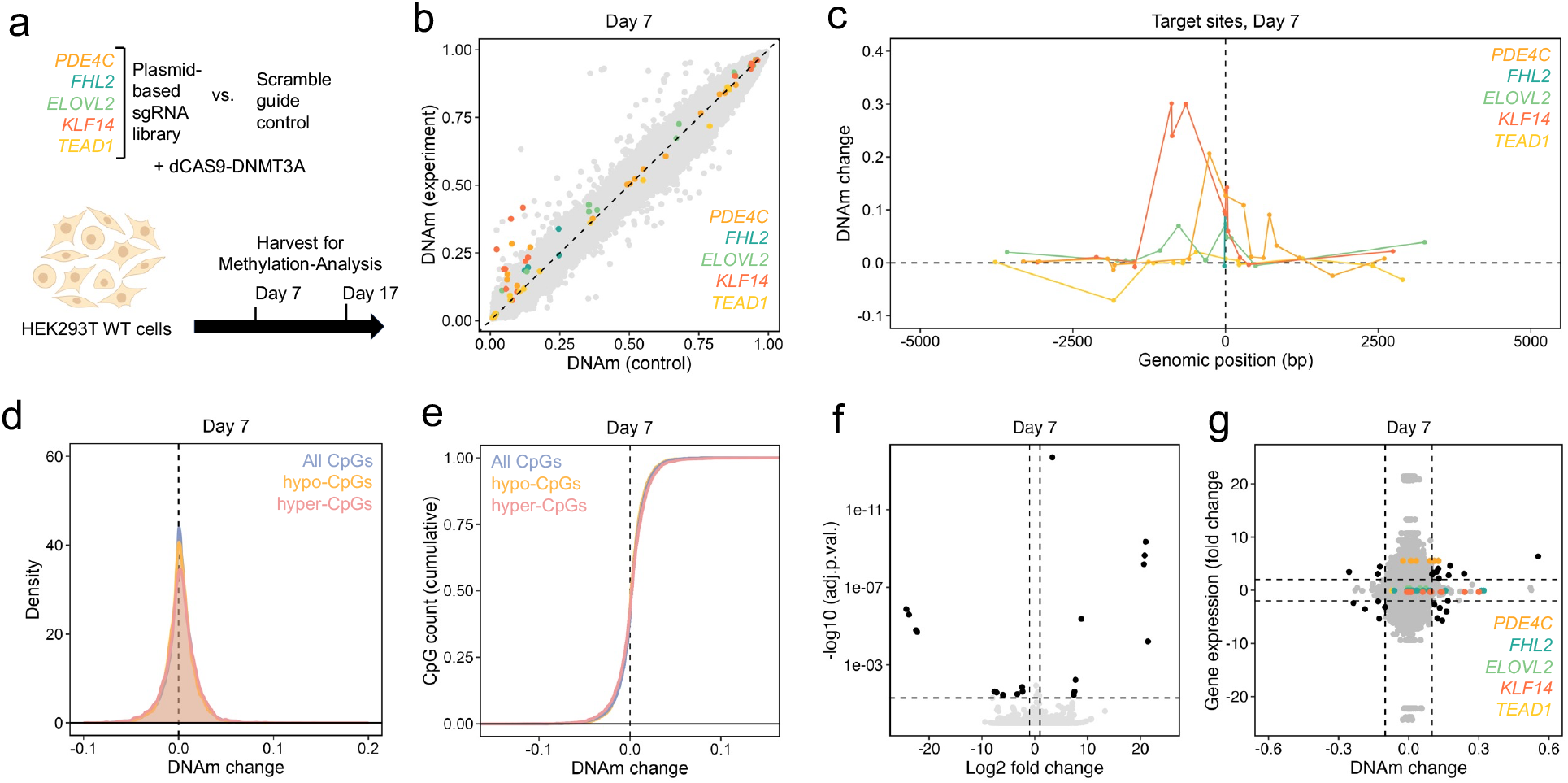
Multiplexed epigenetic editing at age-hypermethylated genomic regions. **a)**Scheme of multiplexed epigenetic editing at five genomic regions that gain DNAm with aging. **b)**Scatterplot of DNAm changes across the three replica. CpGs corresponding to the targeted genes are highlighted in the corresponding color. **c)**Genomic tracks of differential DNAm in a 10k base window centered at the corresponding target site (dashed vertical line) in the five targeted genes. **d**,**e)** Gaussian kernel density estimate and cumulative distribution function comparing the entirety of CpGs from the BeadChip with age-associated CpGs (4389 age-hypo and 5328 age-hypermethylated CpGs). In these experiments hardly any reproducible bystander effects were observed. **f)**Transcriptomic changes upon multiplexed epigenetic editing at the age-hypermethylated CpGs. 10 genes were significantly up- and 11 genes significantly down-regulated (adjusted P < 0.05; >2 fold change). **g)**Correlation of differential gene expression with differential methylation. Gene expression and transcripts were matched by gene IDs. Pairs with >2-fold gene expression change and >10% DNAm change are highlighted.

Transcriptomic analysis of these samples by RNA-sequencing revealed moderate but significant up- and down-regulation of 10 and 11 genes, respectively (n = 3; adjusted P < 0.05, Figure 4f). Overall, there was no clear correlation between gene expression changes and DNAm changes (Figure 4g). However, the target gene *PDE4C* was 5.5-fold higher expressed upon multiplexed epigenome editing.

### Resetting DNA methylation at CpGs that become hypomethylated with age

With regard to the possibility of epigenetic rejuvenation, we next attempted to manipulate genomic sites that lose DNAm with age. To this end, we multiplexed guide RNAs for *COL1A1, AKAP8L, CSNK1D, MEIS1-AS3* and *IGSF11* ^*22*,*28*^, as compared to scramble guide RNAs for control (n = 3; Figure 5a). Initial experiments indicated that modulation of CpGs that become hypomethylated with aging might be less stable and therefore we analyzed DNAm profiles already after three days. Many CpGs gained methylation including CpGs at the target regions of *AKAP8L, CSNK1D*, and *MEIS1-AS3* (Figure 5b,c). These changes were also distributed in a 1.5 kb area around the target regions (Figure S5a). However, after 15 days these targeted gains of DNAm were already lost (n = 3; Figure 5d). Thus, DNAm gains at age-associated hypomethylated regions appear to be less stable than at the hypermethylated CpGs.

**Figure 5:**
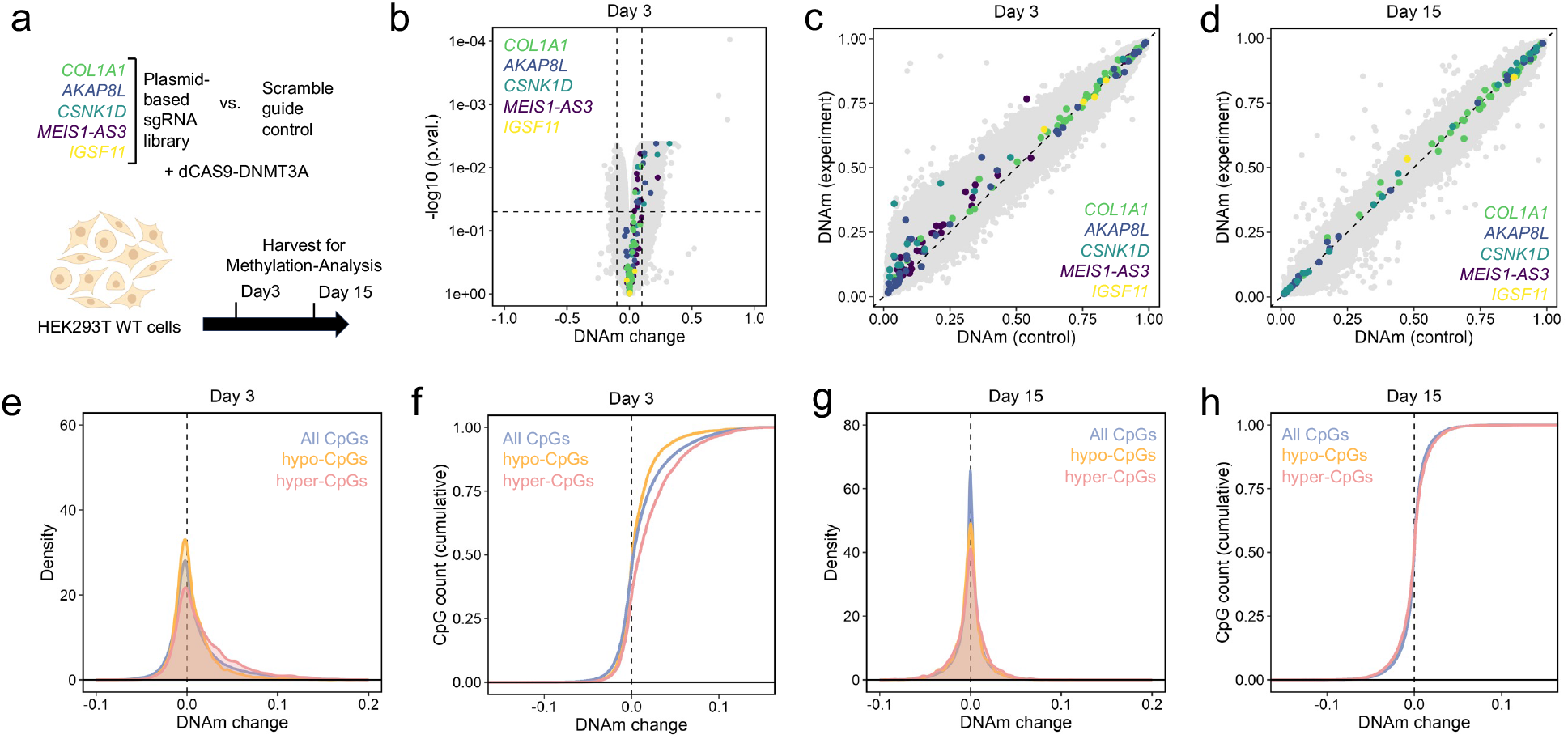
Multiplexed editing at age-hypomethylated regions. **a)** Scheme of multiplexed epigenetic editing at five genomic regions that lose DNAm with aging. **b**,**c)** Three days after transfection, DNAm changes were analyzed by a vulcano plot (b) and a scatter plot (c). CpGs corresponding to the five target genes are highlighted by the corresponding colors (n = 3). **d)**At 15 days after transfection, the gains in DNAm in the target regions were hardly observed anymore. **e)**At day three after transfection, gaussian kernel density estimate at age-associated CpGs (4389 age-hypo and 5328 age-hypermethylated CpGs) showed that bystander modifications were enriched at genomic regions that gain methylation with age. **f)**Cumulative distribution of DNAm at age-associated CpGs at day three after transfection. Notably, bystander effects are underrepresented at age-hypo and overrepresented at age-hypermethylated CpGs. **g**,**h)** At day 15 after transfection kernel density estimate and cumulative distribution did not reveal bystander effects anymore.

At day three, there were 19,254 significant hypermethylated and 88 hypomethylated bystander effects (abs. diff. beta-value > 0.1 and p-value < 0.05). These bystander modifications were again not enriched in predicted off-target regions of the guide RNAs for the five hypomethylated regions (Figure S5b). We have further analyzed whether the bystander effects observed in the initial *PDE4C* experiment correlated with the bystander effects of hypomethylated CpGs. While some of the bystander modifications were consistent in both experiments, there were also several changes that are exclusively observed targeting the five age-hypomethylated regions, further substantiating the notion that the bystander modifications are not only footprints that are attributed to binding preferences of DNA methyltransferases domains (Figure S5c).

Subsequently, we tested if the day 3 bystander modifications of hypomethylated target regions would enrich for age-associated CpGs. Strikingly, the 5328 age-hypermethylated CpGs were again enriched in these bystanders, whereas the overlap with the 4389 age-hypomethylated CpGs was significantly lower than expected (Figure 5e,f). This was somewhat surprising, since we have now modified CpGs that become hypomethylated with aging, but it clearly demonstrated that age-associated DNAm is coherently mediated, particularly at age-hypermethylated CpGs. However, at day 15 there were no significant bystander modifications anymore, and thus, also age-associated CpGs did not reveal consistent modifications (Figure 5g,h).

### Epigenetic editing accelerates epigenetic clocks in T cells

Since our age-associated CpGs were identified in blood, we have next aimed for epigenetic modulation in primary T cells. We have again co-transfected the five guide RNAs that target age-associated hypermethylated CpGs (*PDE4C, FLH2, ELOVL2, KLF14*, and *TEAD1*; n = 2; controls with scramble guide RNA and dCas9-DNMT3A, Figure 6a). In fact, after 21 days we observed site specific DNAm, particularly at the target sites of *FHL2* and *KLF14* (Figure 6b). In these experiments we observed many bystander modifications, but none of them were significant due to variation between the two biological replicas. Notably, hypo-as well as hyper-methylated bystander modifications were enriched at CpGs that gain or lose-DNAm with aging in blood, and this was particularly observed at age-hypermethylated CpGs (Figure 6c,d). Thus, the induction of DNAm at CpGs that gain DNAm with age did not have complementary effects on age-hypo and age-hypermethylated CpGs, but the bystander effects were clearly enriched at these sites.

**Figure 6:**
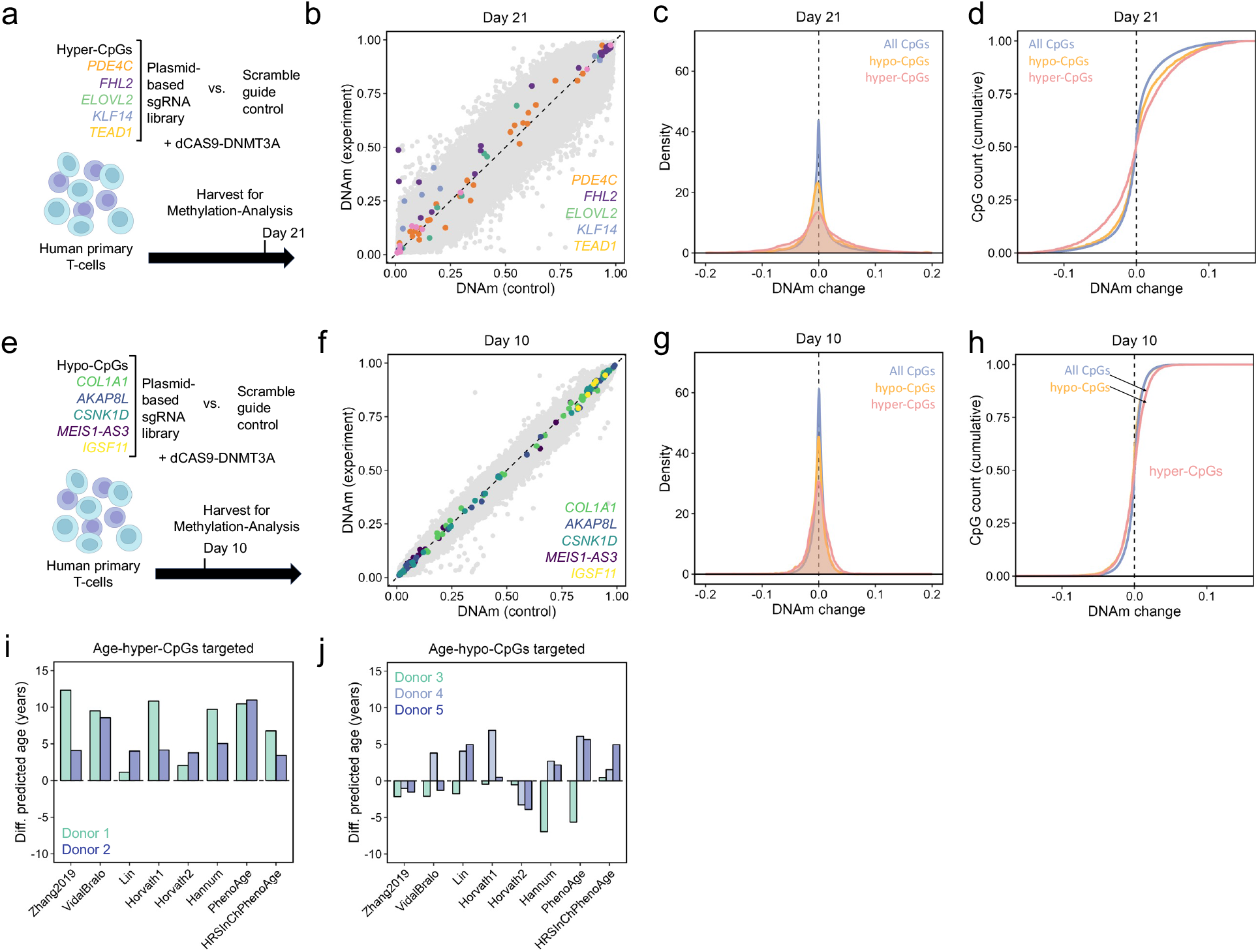
CIRSPR-epigenetic editing in human primary T-cells. **a)**Scheme of multiplexed epigenetic editing at five genomic regions that gain DNAm with aging. **b)**Scatterplot of Illumina BeadChip data showing clear methylation changes at *FHL2* and *KLF14* (n = 2). **c, d)** Gaussian kernel density estimate and cumulative distribution function showing significantly different methylation profiles at hyper- and hypo-CpGs (asymptotic two-sided Kolmogorow-Smirnov test P < 10^-15^). **e)**Scheme of multiplexed epigenetic editing at five genomic regions that lose DNAm with aging. **f)**Scatterplot of Illumina BeadChip data revealing no methylation change after 10 days (n = 3). **g, h)** Gaussian kernel density estimate and cumulative distribution function. The genome-wide epigenetic landscape is less affected when targeting age-hypomethylated CpGs (10 days of culture). **i, j)** Estimation of epigenetic age with eight different epigenetic clocks: : Zhang ^29^, Vidal-Bralo ^30^, Lin ^4^, Horvath 1 (multitissue) ^9^, Horvath 2 (skin and blood) ^31^, Hannum ^28^, and PhenoAge and updated PhenoAge ^7^. Bar plots depict the deviation of epigenetic age predictions upon targeted modification at the age-hypermethylated (i) and the age-hypomethylated CpGs (j), as compared to the scramble guide RNA controls (difference in years).

Analogously, we next targeted DNAm at the five age-hypomethylated CpGs at *COL1A1, AKAP8L, CSNK1D, MEIS1-AS3*, and *IGSF11* (n = 3, Figure 6e). After 10 days, we did not observe clear DNAm changes in the genome wide DNAm profiles at the target regions, and there were only very moderate bystander modifications (Figure 6f). These results further support the notion that epigenetic engineering was more stable at the age-hyper CpGs than at the age-hypo CpGs. Accordingly, the genome-wide aging signature was less affected with transient manipulations of age-hypomethylated CpGs – but there were still moderate enrichments (Figure 6g,h).

As both epigenetic engineering-approaches at either age-hyper or age-hypomethylated CpGs were clearly enriched at age-associated genomic regions, we have subsequently analyzed how this might impact epigenetic clocks. To this end, we have utilized eight different epigenetic algorithms that have been trained for blood samples: Zhang ^29^, Vidal-Bralo ^30^, Lin ^4^, Horvath 1 (multitissue) ^9^, Horvath 2 (skin and blood) ^31^, Hannum ^28^, PhenoAge ^7^, and updated PhenoAge ^7^. In fact, all of these epigenetic clocks revealed an epigenetic age-acceleration of up to ten years after epigenetic editing at the age-hypermethylated regions in comparison to the un-targeted dCas9-DNMT3A (Figure 6g). In contrast, the transient effects at age-hypomethylated regions had less impact on epigenetic clocks and resulted in a moderate acceleration of epigenetic clocks in two donors (Figure 6h). Overall, these results indicate that it is possible to modulate epigenetic clocks by epigenome editing – but so far only to accelerate these epigenetic modifications.

## Discussion

It is largely unclear how DNA methyltransferases are guided to specific sites in the genome. The enormous complexity of these genome-wide epigenetic modifications suggests that they might be controlled by some kind of interactive network, possibly involving other epigenetic features, such as the histone code and chromatin architecture. The results of this study demonstrate that targeted epigenetic editing not only unravels the relevance of site-specific DNAm patterns at the target site, but also opens new opportunities to elucidate mechanisms that govern genome wide epigenetic interactions.

It was unexpected that targeted gains of DNAm can be stable for more than three months. Previous studies have only described transient impact of DNMT3A-mediated epigenome editing ^13,14^. In fact, at genomic regions that become hypomethylated with age we only observed transient gains in DNAm. It is conceivable that epigenetic clock sites that gain DNAm with age are more susceptible to accumulate DNAm. This may also be the reason why they become consistently methylated with age.

Furthermore, we did not anticipate epigenetic editing to stochastically modulate DNAm at neighboring CpGs on the same DNA strand. We assumed that successful targeting of the methyltransferases to a specific genomic region would result in homogeneous methylation at that region, but bisulfite amplicon sequencing demonstrated otherwise. We have previously shown that also during aging age-associated DNAm is not homogeneously acquired at neighboring CpGs ^22^. In fact, recent reports demonstrate that large parts of the predictive accuracy of epigenetic clocks can be explained by stochastic processes ^32,33^. However, it remains unclear why this stochastic drift occurs preferentially at specific age-associated CpGs and why stochastic processes seem to contribute less to clock sites that rather reflect biological age than chronological age. It is conceivable that DNAm patterns are changing in a much more dynamic manner towards a region-specific equilibrium than generally anticipated. This might explain why we even observed a gain in DNAm over time at some CpGs within the *PDE4C* amplicon, resulting in a more homogeneous pattern at neighboring CpGs. On the other hand, due to the incoherent modifications at neighboring CpGs the fraction of modified DNA strands may be underestimated by analysis of DNAm levels at individual CpGs. Other authors have described successful modification of about 50% of the DNA strands ^34^ and this was in line with our results based on Illumina BeadChips or pyrosequencing. However, the fraction of modified reads indicated that up to 61% of the DNA alleles were stably modified. In fact, analysis at day three revealed that initially even 88.9 % of the DNA allels were successfully modified, demonstrating that epigenetic editing was highly efficient in the large majority of the cells.

Off-target effects have been described as a major problem for epigenome editing ^35^. The footprints of CRISPR-guided DNAm approaches have been analyzed in detail before ^17^ and it has been suggested that this is particularly based on the utilized targeting tool ^23^. Notably, some of these off-targets were even hypomethylated, particularly at repetitive sequences, and the majority of the off-targets was not associated with sequence homologies of gRNAs ^17,23^. There have been attempts to decrease off-target effects, e.g. by different methyltransferases or targeting constructs ^36^. However, the findings of our current study indicate that the off-target effects might rather reflect controlled epigenetic changes with a network: 1) These DNAm are observed in a highly reproducible manner in independent biological replica, 2) they are observed in a very similar manner with different CRISPR constructs, 3) they include significant gains and losses of DNAm, 4) footprinting was also observed when we normalized the experiments for controls with epigenome editors and scrambled guide RNA, 5) bystander effects were not related to sequence homology of the guide RNAs, and 6) there was a low correlation of bystander modifications upon targeting age-hyper or age-hypomethylated CpGs. Therefore, we suggest that these modifications should rather be termed bystander modifications than off-target effects.

So far, it is unclear how bystander modifications might be regulated. Our 4C-sequencing results indicate there might be chromatin interaction between genomic sites that have coherent bystander modifications. However, this does not fully explain all of the complex DNAm changes – and it is as well also unclear how the interaction in chromatin conformation is governed. Architectural proteins, such as CCCTC-binding factor (CTCF), can medicate intra- and interchromosomal interactions. We previously demonstrated that age-associated hypermethylation indeed peaks near CTCF binding sites ^22^. Knockdown of CTCF did not interfere with epigenetic clocks ^11^, but it is still conceivable that the formation of chromatin loops and interactions contributes to these bystander modifications.

Perhaps the most striking finding of this study was that the bystander modifications of age-associated regions were significantly enriched at other age-associated CpGs throughout the genome – particularly at CpGs that gain DNAm with aging. Hypermethylation with aging occurs preferentially at regions with high CpG density ^37^, which may even be sufficient to predict aging trajectories ^38^. In fact, bystander modifications were highly enriched next to CpG islands too. Previous work identified that CpGs in a low CpG context with flank nucleotides A and T are prone to lose methylation with age (hypo-CpGs or solo-WCGW’s)^26^, which was also observed in our analysis. In contrast, our bystander modifications had rather opposite flank nucleotides. Thus, bystander modifications can not only be attributed to the above mentioned site-specific characteristics, further pointing towards an epigenetic network.

Our results indicate that it is possible to interfere with epigenetic clocks by epigenetic editing. So far, our stable epigenetic editing at age-hypermethylated CpGs only accelerated epigenetic age in T cells. Increasing epigenetic age may not appear attractive as compared to the perspective of rejuvenation, but there are also applications were increasing epigenetic age might be useful. For example, iPSCs are often used as a model system to study age-associated diseases – since these cells become epigenetically rejuvenated during reprogramming into pluripotent state, epigenetic editing with accelerated age-associated DNAm may better reflect the aging phenotype.

Taken together, our results indicate that epigenetic editing at age-associated regions can perturb an epigenetic network and evoke coordinated epigenetic bystander modifications. These modifications are enriched at other age-associated CpGs and therefore have an impact on epigenetic clocks. While many questions remain unanswered, our findings open the perspective to control epigenetic aging in the future and thereby answer the question of whether the epigenetic clocks are *per se* functionally relevant.

## Methods

### Constructs for epigenome editing

We use two constructs for epigenome editing: 1) dCAS9-DNMT3A/3L co-expressed with enhanced green fluorescent protein (eGFP) for selection ^12^ (Addgene, Catalogue number 128424), and 2) CRISPRoff that comprises the DNMT3A/3L domain at the N-terminus, and blue fluorescent protein (tagBFP) with a KRAB domain at the C-terminus (Addgene 167981; Figure 1A)^14^. Guide RNAs were ordered as dsDNA from Integrated DNA Technologies (Supplemental Table S1) and cloned into the expression vector (Addgene 44248) by double digest with BstXI and NotI (Thermo Fischer Science). Plasmids were amplified in DH5-alpha competent cells (EC0112, Thermo Fisher), isolated using a kit (M+N Nucleospin Plasmid kit). Successful cloning was confirmed by enzymatic restrition and Sanger sequencing (Eurofins).

### Cell culture

HEK293T cells were cultured in Dulbecco’s Modified Eagle Medium (Gibco™) with 10% fetal calf serum (FCS; Bio&Sell) and Penicillin-Streptomycin (Life Technologies) at 37°C / 5% CO_2_, and passaged manually at 80 - 90% confluency. The identity of this cell line was reconfirmed by their DNAm profiles (Illumina BeadChip).

Primary human T cells were isolated from peripheral blood of blood donors at the Clinic of Transfusion Medicine, RWTH University Hospital Aachen. Samples were taken after written and informed consent following the procedure approved by the ethics committee of RWTH Aachen University (EK 206/09) and supported by the RWTH central biomaterial bank (cBMB). Peripheral blood mononuclear cells (PBMCs) were harvested and T cells were isolated by magnetic separation with the human Pan T Cell isolation kit and stimulated using TransAct (both Miltenyi Biotec). Cells were then cultured in TexMacs medium (Miltenyi Biotec) with 10% FCS (Bio&Sell), IL-2 (50 IU/mL, Gibco^TM^) and Penicillin Streptomycin 100U/ml (Gibco^TM^) at 37°C / 5% CO_2_.

### Transfection and selection of cells

HEK293T cells were transfected with the TransIT®-LT1 transfection reagent (Mirus Bio) in 6-well plates with 1.25μg of plasmids for the CRISPR-construct and 1.25μg of plasmids for single gRNA. We either pooled two gRNAs targeting *PDE4C* or up to nine gRNAs for multiplexed editing. After 24 hours we selected the transfected cells either by puromycin treatment (2 ug/ml for 2 days) or flowcytometric sorting for double positive cells (BD FACSAria^TM^ fusion).

Transfection of T cells was performed using the NEON transfection system (Invitrogen). 5 million T cells were transfected with 5µg of plasmids for CRISPR-constructs and 5μg of gRNA plasmids inside of a NEON 100ul pipet tip with 1 pulse (20ms) at 2100V. After 24 hours, cells were resuspended in MACS buffer for flowcytometric sorting for double positive cells (BD FACSAria^TM^ fusion) and further cultured in TexMACS medium as indicated above for up to 21 days.

### Analysis of DNA methylation profiles

Genomic DNA was isolated from HEK293T or primary T cells using the Macherey Nagel Tissue kit. DNA was bisulfite converted and hybridized to EPIC Illumina BeadChips (either version 1 or 2) at Life and Brain GmbH, Bonn, Germany. Initial quality control was performed on IDAT files using the minfi package (version 1.48.0)^39^ and for subsequent preprocessing we used the sesame package (version 1.20.0): Data was normalized with noob and probes with a detection p-value > 0.05 in at least one sample were removed^40,41^. Furthermore, we excluded probes associated to X or Y chromosomes, SNPs, and CpGs flagged in the b5-manifest (Illumina). Differential methylation analysis was performed using limma. Probes with an p-value < 0.05 (Benjamini Hochberg adjusted) and at least 10% mean difference in DNAm were considered significantly differentially methylated. Genomic feature annotation and flank sequences (hg38 genome) was obtained from the manifest file EPICv2 (Illumina). Enrichment of CpGs in genomic features was analysed using Chi^2^ tests. Significance between mean DNAm changes of subgroups was estimated by unpaired t-tests. All plots were generated in R using ggplot2 and ggforce.

### Pyrosequencing

Genomic DNA was bisulfite converted with the EZ-DNA methylation kit (Zymo) and PCR amplified at the age-associated regions with the PyroMark PCR kit (Qiagen). 10μl of PCR-product and 2µl of sequencing primer (4μM) were then pyrosequenced on a Pyromark Q48 Autoprep (Qiagen), as described before ^22^. Primer sequences are provided in Supplemental Table S2.

### Bisulfite barcoded amplicon sequencing

Bisulfite barcoded amplicon sequencing at age-associated genomic regions was performed as described in detail before ^22^. In brief, bisulfite-convered genomic DNA was amplified with the PyroMark PCR kit (35 cycles; Qiagen) with primers containing handle sequences. The amplicons were purified with the Select-A-Size DNA Clean & Concentrator Magbead kit (Zymo) and further amplified with a second PCR (10 cycles) to add barcodes, i5 and i7 adapters. The PCR products were pooled equimolarly and washed in a size-selection column (Zymo Select-A-Size DNA Clean & Concentrator). Sequencing was performed using the Illumina v2 Nano reagents and flow cell kit with 50% PhiX control on MiSeq (250bp, paired end). PCR primers are provided in Supplemental Table S3. We used Bismark to obtain methylation calls ^42^ and excluded reads with at least one undefined methylation status. Results were visualized using ggplot in R or matplotlib and seaborn packages in python.

### Chromatin conformation capture 4C analysis

Preparation of the i4C templates was performed as previously described ^43^. In brief, 10^7^ HEK293T cells were used for isolation of nuclei in a near-physiological isotonic buffer. The primary restriction enzyme digestion was performed with 800 U *Nla*III or *Apo*I (New England Biolabs, R0125 and R3566). Following *in situ* ligation of digested chromatin, nuclei were lysed and the isolated DNA was digested with *Cvi*QI (New England Biolabs, R0639), circularized by ligation and purified. The library was amplified by inverse PCR with viewpoints-specific primers (Supplemental Table S4) coupled to Illumina adapters for sequencing. Following inverse PCR, the i5 and i7 Illumina adapters were used to generate NGS-compatible libraries as previously described ^44^. Amplicons were sequenced in a 50 bp paired-end mode on the Illumina NextSeq2000 platform (Genomics facility, Erasmus MC Rotterdam). Viewpoint primers sequences were trimmed using cutadapt, and the remaining sequences were aligned to the reference genome (hg38) by HISAT2. After quantification of reads per fragement, 4C data was overlapped with the positions of the Illumina Beadchip (Illumina manifest file, hg38) using the GenomicRanges package in R. We further analysed the association of 4C coverage and aging positions or bystander positions using Chi2 testing and logistic regression.

### Gene expression analysis

RNA was isolated using the RNA plus kit (Macherey Nagel). Library preparation (QuantSeq 3’mRNA library prep, Lexogen) and sequencing (Novaseq 6000, 100bp single end) was performed at Life and Brain GmbH. Preprocessing of fastq files was carried out using the nextflow-core pipeline: Adapters and low quality reads were removed using Trim Galore. We used STAR for alignment (hg38 genome) and counts matrices were generated by Salmon. Counts matrices were normalized and differential gene expression was performed using DESeq2 in R ^45^. P values were calculated using a negative binomial GLM function and Wald-test. Differential gene expression as considered significant with a p value smaller than 0.05 (Benjamini Hochberg adjusted) and at least two-fold change. For integration of transcriptomic changes and DNAm changes, BeadChip probes were matched to transcripts based on Gene ID.

## Additional information

### Author Contributions

S.L., M.M., A.P. and W.W. contributed to the experimental design. S.L., J.I., and A.M. carried out the experiments. S.L., M.M., and M.V.B. performed the bioinformatic analysis. M.W. contributed viable materials. W.W. initiated and supervised the study. S.L. wrote the initial draft of the manuscript. All authors revised and approved the final version of the manuscript.

## Acknowledgements

This work was supported by the Flow Cytometry Facility and the Genomics Facility, both core facilities of the Interdisciplinary Center for Clinical Research (IZKF) Aachen within the Faculty of Medicine at RWTH Aachen University. We acknowledge the Genomics Core Facility at Erasmus Medical Center, Rotterdam, Netherlands for performing the sequencing and preprocessing pipeline for 4C-analysis.

