## Supplemental Figures and Tables for "Epigenetic editing at individual age-associated CpGs affects the genome-wide epigenetic aging landscape"

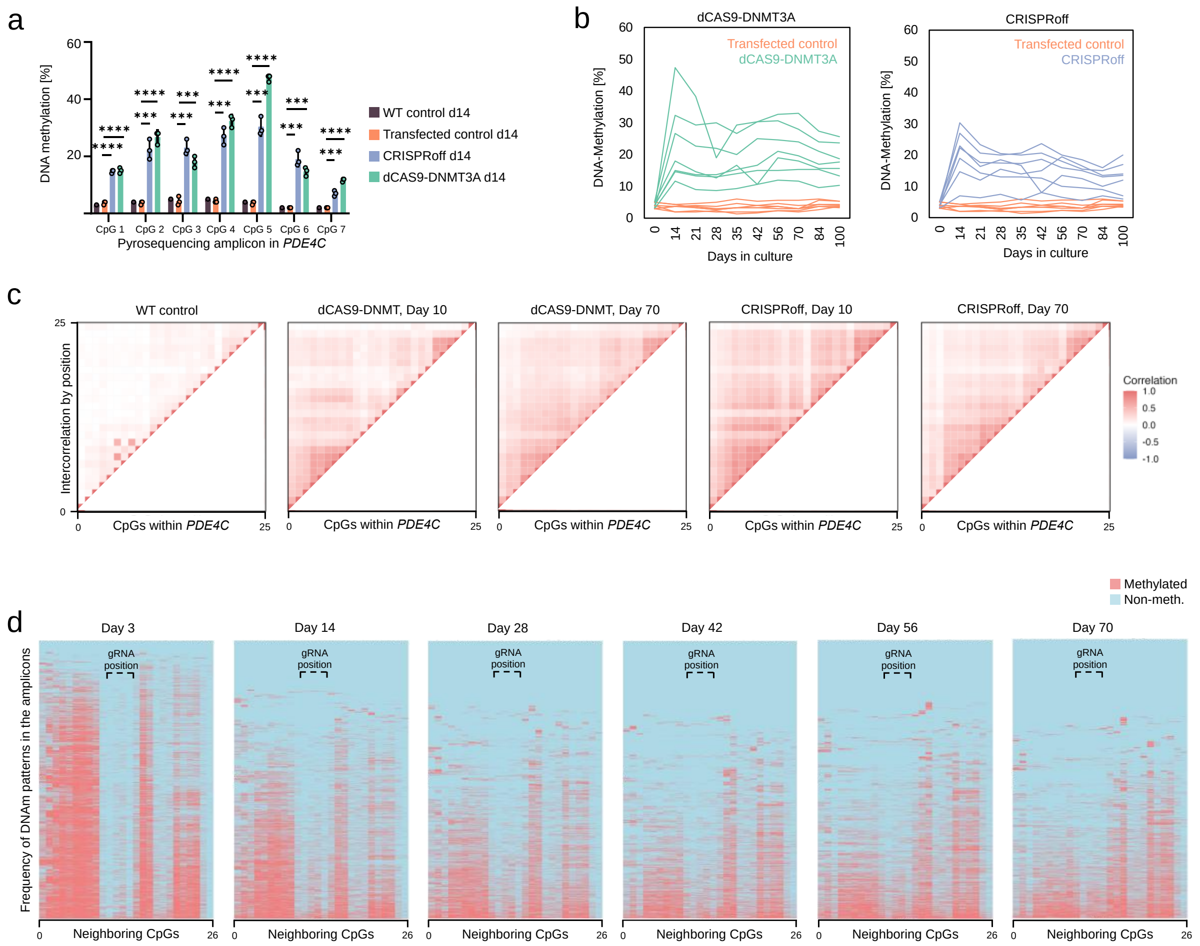


### Supplemental Figure S1: DNA methylation in amplicons of the target region.

**a)** Pyrosequencing measurement of HEK293T cells after epigenetic modification at *PDE4C* (three replicates). DNAm levels at the seven CpGs in the pyrosequencing amplicon are depicted at day 14 (***P < 0.001; **** P < 0.00001).

**b)** Pyrosequencing of the same samples over a time course of up to 100 days confirmed that the modified DNAm is stable at this site.

**c)** To better understand if neighboring CpGs on the same DNA strand become coherently modified by CRISPR-guided epigenetic editing, the correlation of DNAm was analyzed at the 26 neighboring CpGs within the bisulfite barcoded amplicion sequencing reads. The heatmap depicts Pearson correlations between individual CpGs.

**d)** Heatmaps showing the complete time-course experiment upon modification with dCAS9-DNMT3A, corresponding to Figure 1e.

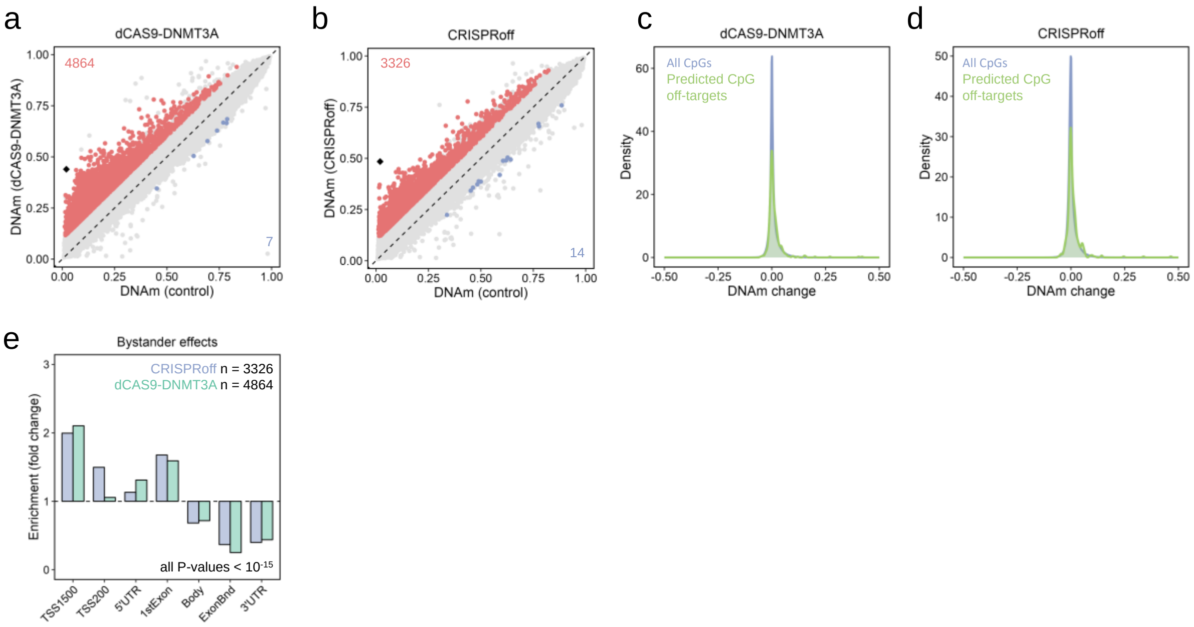


**Supplemental Figure S2: Further analysis of epigenetic bystander modifications.**

**a,b)** Scatterplots of Illumina BeadChip data demonstrate DNAm changes in HEK293T cells upon targeted methylation at *PDE4C* with either dCas9-DNMT3A (a; n = 3), or CRISPRoff (b; n = 3). Significant CpGs are highlighted in analogy to Figure 2a,b.

**c,d)** Due to sequence homology, 123 genomic regions were identified as potential off-target binding sites. Within a window of 5kb, we identified 300 CpGs in these regions. Predicted off-target binding sites for *PDE4C* gRNAs did not reveal clear enrichment in bystander modifications of dCas9-DNMT3A (c), or CRISPRoff (d).

**e)** Enrichment of the bystander effects in relation genomic regions: TSS1500: 1500 bp upstream of transcription start site (TSS); TSS200: 200 bp upstream of TSS; UTR: untranslated region. Enrichment was calculated in relation to all CpGs on the array and was highly significant for all categories (Chi$^2$ test).

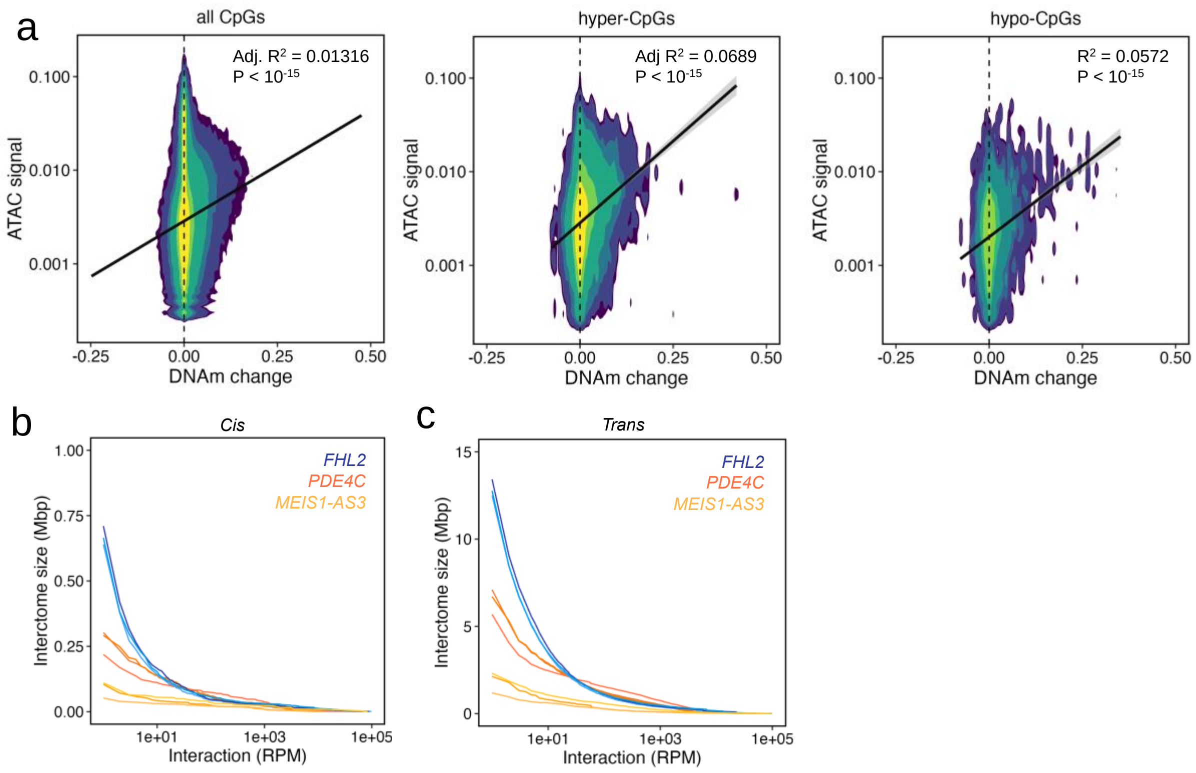


**Supplemental Figure S3: Further analysis of ATAC-seq and 4C-sequencing data.**

**a)** Multivariate kernel density estimate of ATAC-seq and DNA methylation changes (dCAS9-DNMT3A). While differentially methylated positions exhibited in tendency higher chromatin accessibility the overall correlation was rather low. In analogy, we analyzed the 4389 age-hypo and 5328 age-hypermethylated CpGs.

**b)** Cis-interactions (on the same chromosome) of 4C-Sequencing with around 20 million reads per sample. Particularly the two viewpoints at positively age-correlated regions in *PDE4C* and *FHL2* exhibited cis-interactions spanning a mean of 271 and 557 kbp, respectively. In contrast, the 4C reads of age-hypomethylated region in *MEIS1-AS3* comprised only 89 kbp.

**c)** Trans-interactions (across different chromosomes) of 4C-sequencing. The viewpoints at positively age-correlated regions like *PDE4C* and *FHL2* exhibited trans-interactions spanning a mean of 6.5 and 13.0 mbp, respectively.

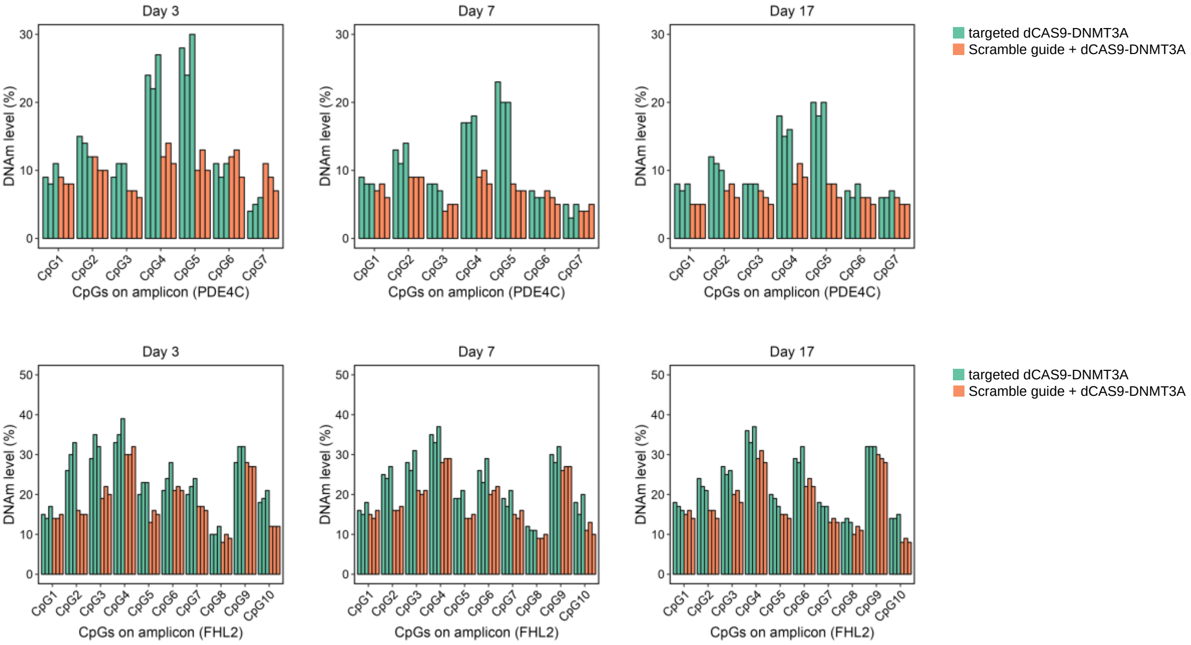


**Supplemental Figure S4: Pyrosequencing validation of targeted epigenetic modification.**

Pyrosequencing measurements of demonstrating stable DNAm levels at *PDE4C* and *FHL2* (triplicates). HEK293T cells were either transfected guides for *PDE4C, FLH2, ELOVL2, KLF14,* and *TEAD1* together with dCas9-DNMT3A, or scramble guide RNA and dCas9-DNMT3A. The epigenetic modifications both regions were stable for at least 17 days. DNAm was significantly higher in experimental conditions across all timepoints (p-value < 0.05, unpaired t-test).

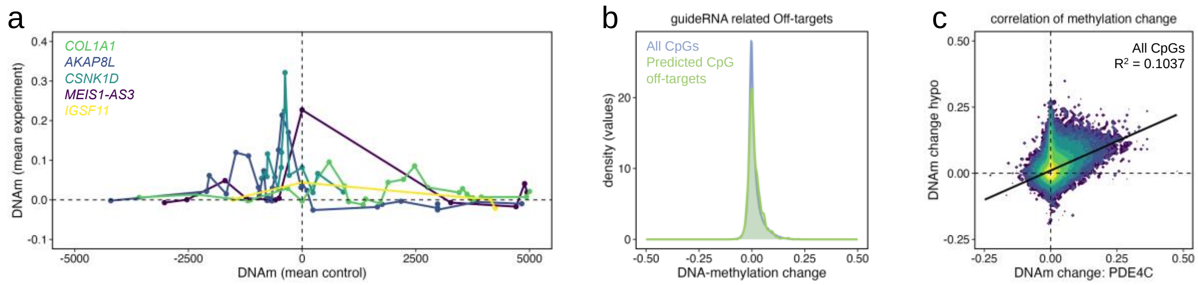


**Supplemental Figure S5: Targeted modifications at age-hypomethylated regions.**

**a)** Genomic track of DNAm changes at fife hypo-CpGs analogue to Figure 4c.

**b)** Kernel density estimate revealing no methylation changes at 692 predicted off-target binding sites (sequence homology) compared to all CpGs on BeadChip.

**c)** Multivariate kernel density estimate and linear regression comparing DNAm changes with epigenome editing between *PDE4C* and hypo-CpGs.

**Table S1: Guide RNA sequences for age-associated hyper-CpGs**

| Target gene and no. | sgRNA-guide sequence | Target CpG ID (Illumina BeadChip) |
|---|---|---|
| Hypermethylated with age: | | |
| *PDE4C* #1 | 5‘-TCTGGCGCATGGAGAACCTG-3‘ | cg17861230 |
| *PDE4C* #2 | 5‘-TCGGATCCGGACAAGTCCGC-3‘ | cg17861230 |
| *ELOVL2* #1 | 5‘-GACGAAACGGCCCCGAGGCT-3‘ | cg16867657, cg21572722 |
| *ELOVL2* #2 | 5‘-GGGGAGAAGCAGTATCGTGC-3‘ | cg16867657, cg21572722 |
| *FHL2* #1 | 5‘-GAGAAGCCCCAGGACGTGCCG-3‘ | cg22454769, cg06639320 |
| *FHL2* #2 | 5‘-GACTGTGCTCCCAAGACCCGG-3‘ | cg22454769, cg06639320 |
| *KLF14* #1 | 5‘-GCAACCCAGAAGTTCCGACTG-3‘ | cg22285878 |
| *KLF14* #2 | 5‘-GACGGAGCACGGGATCGGGT-3‘ | cg22285878 |
| *TEAD1* #1 | 5‘-GCACAAGTCTGCGCAGACCC-3‘ | cg04940570 |
| Hypomethylated with age: | | |
| *MEIS1-AS3* #1 | 5‘-CAAGTGAATAACTACTCAGC-3‘ | cg11807280 |
| *COL1A1* #1 | 5‘-TAAGATTGGAGAAGGTTGAC-3‘ | cg18618815 |
| *COL1A1* #2 | 5‘-TCTCCTATAGAAGAACTCCC-3‘ | cg18618815 |
| *AKAP8L* #1 | 5‘-CACACGTGTTAAGTACTCTT-3‘ | cg25533247 |
| *CSNK1D* #1 | 5‘-GCCTTACCTCCTGACACCTA-3‘ | cg19761273 |
| *IGSF11* #1 | 5‘-TAATCAATAAGACCTTCCTA -3‘ | cg00329615 |
| *IGSF11* #2 | 5‘-GGCTACGATTCAGGGACACC-3‘ | cg00329615 |

**Supplemental table S2: Primers for pyrosequencing**

| Gene name | Label | Primer sequence |
|---|---|---|
| *PDE4C* | Forward | 5‘-AGGTTTGTAGTAGGTTGAG-3‘; |
| | Reverse | Biotin-5‘-AACTCAAATCCCTCTC-3‘ |
| | Seq-Primer | 5‘-GTTATAGTATGATTAGAGTTT-3‘ |
| FHL2 | Forward | 5‘-GTGTTTTTAGGGTTTTGGGAGTATAGTAGT-3‘ |
| | Reverse | Biotin-5‘-CACCTCCTAAAACTTCTCCAATCTCC-3‘ |
| | Seq-Primer | 5‘-GGTTTTGGGAGTATAGTAGTT-3‘ |
| *ELOVL2* | Forward | 5‘-Biotin-GGGAGGGGAGTAGGGTAAGTGA-3‘ |
| | Reverse | 5‘-CCATCTAAACAACCAATAAATATTCCTAAA-3‘ |
| | Seq-Primer | 5‘-AATAAATATTCCTAAAACTC-3‘ |
| *MEIS1-AS3* | Forward | 5‘-ATTTTTGTTTGAAGGTTTTTATAAAATATG-3‘ |
| | Reverse | 5‘-ACCTTTAAAACAACAAAATAAATCACACT-3‘ |
| | Seq-Primer | 5‘-ACCATACTTAACATCCA-3‘ |
| *AKAP8L* | Forward | 5‘-TGAAGTTTGGAATTTATGATTTGTTTAAG-3‘ |
| | Reverse | Biotin-5‘-CCCAAAAAACAAACTAAAAACATAACTAAT-3‘ |
| | Seq-Primer | 5‘-GAGAGATTTTGTAATAGTGTA-3‘ |
| *CSNK1D* | Forward | 5‘-GGAGGTTTTGATGTTTAGTTTGAAGAT-3‘ |
| | Reverse | Biotin-5‘-CAAATCCAACACAAATAAAAATATTAACTC-3‘ |
| | Seq-Primer | 5‘-GGTTAGATTATTTGTTTTTTTTTAG-3‘ |
| *COL1A1* | Forward | 5‘-TTGAAGGGAAGAGGTAAGGAAGATTTTA-3‘ |
| | Reverse | Biotin-5‘-TAACCCATCTTTTTCCTTCTTCTCA-3‘ |
| | Seq-Primer | 5‘-AATTTGTATAGAGAGTGTTTATTG-3‘ |
| *IGSF11* | Forward | 5‘-GTTGGATAGTTTGTGGGTAGAAAATTTA-3‘ |
| | Reverse | Biotin-5‘-ATTATTCATTCATTATTCTCCTTAAAAAAATCTTATT-3‘ |
| | Seq-Primer | 5‘-AGAAGTTAAGAAGGTATAGATA-3‘ |

**Supplemental table S3: Primers for bisulfite amplicon sequencing**

| Gene Name | Label | Primer sequence |
|---|---|---|
| PDE4C, 1st PCR | Forward | 5‘-CTCTTTCCCTACACGACGCTCTTCCGATCTTATGGAGAATTT GGGG-3‘ |
| | Reverse | 5‘-CTGGAGTTCAGACGTGTGCTCTTCCGATCTCTACAAAAAAC CCCTACC-3‘ |
| Adapters, 2nd PCR | i5 | 5‘-AATGATACGGCGACCACCGAGATCTACACTCTTTCCCTACAC GACGCTCTTCCGATCT-3‘ |
| | i7 | 5‘-AGATCGGAAGAGCACACGTCTGAACTCCAGTCAC[Barcode]AT CTCGTATGCCGTCTTCTGCTTG-3‘ |

**Supplemental table S4: Primers for i4C sequencing**

| Primer | Location (hg38) | Adapter primer Illumina; view-point specific sequence |
|---|---|---|
| Pri-PDE4C-NlaIII | chr19:18233097-18233114 | 5'-TCGTCGGCAGCGTCAGATGTGTATAAGAGACAG-3'; 5'-AGGTGCTTCGGGGCTCTG-3'; |
| Sec-PDE4C_CviQI | chr19:18232652-18232672 | 5'-GTCTCGTGGGCTCGGAGATGTGTATAAGAGACAG-3'; 5'-CCAGATGTGTTTGGGGTGCTC-3' |
| Pri-FHL2-NlaIII | chr2:105399436-105399455 | 5'-TCGTCGGCAGCGTCAGATGTGTATAAGAGACAG-3'; 5'-AGGGGGTCACTTCTCAGGAG-3' |
| Sec-FHL2_CviQI | chr2:105399228-105399247 | 5'-GTCTCGTGGGCTCGGAGATGTGTATAAGAGACAG-3'; 5'-AAGAAAGGAGCCCTGGCAAA-3' |
| Pri-MEIS1-ApoI | chr2:66427399-66427383 | 5'-TCGTCGGCAGCGTCAGATGTGTATAAGAGACAG-3'; 5'-GAAGGCTTCCTGCGGCG-3' |
| Sec-MEIS1_CviQI | chr2:66427579-66427598 | 5'-GTCTCGTGGGCTCGGAGATGTGTATAAGAGACAG-3'; 5'-AGTGACTAGAGCACGTTCGC-3' |
